# Surface phenotyping and quantitative proteomics reveal differentially enriched proteins of brain-derived extracellular vesicles in Parkinson’s disease

**DOI:** 10.1101/2022.10.17.512628

**Authors:** Tanina Arab, Yiyao Huang, Rajini Nagaraj, Evan Gizzie, Javier Redding-Ochoa, Juan C. Troncoso, Olga Pletnikova, Tatiana Boronina, Robert N. Cole, Vasiliki Mahairaki, David A. Routenberg, Kenneth W. Witwer

**Affiliations:** Department of Molecular and Comparative Pathobiology, Johns Hopkins University School of Medicine, Baltimore, MD, USA; Meso Scale Diagnostics, LLC, Rockville, MD, USA; Department of Neurology, Johns Hopkins University School of Medicine, Baltimore, MD, USA; Department of Pathology, Johns Hopkins University School of Medicine, Baltimore, MD, USA; Department of Pathology and Anatomical Sciences, Jacobs School of Medicine and Biomedical Sciences, University at Buffalo, Buffalo, NY, USA; Department of Biological Chemistry, Johns Hopkins University School of Medicine, Baltimore, MD, USA; Department of Genetics, Johns Hopkins University School of Medicine, Baltimore, MD, USA; The Richman Family Precision Medicine Center of Excellence in Alzheimer’s Disease, Johns Hopkins University School of Medicine, Baltimore, MD, US

**Keywords:** Parkinson’s disease, microglia, astrocytes, neurons, extracellular vesicles, ectosomes, exosomes, biomarkers, proteomics, progressive supranuclear palsy

## Abstract

Extracellular vesicles (EVs) are produced by all cell types and are found in all tissues and biofluids. EV proteins, nucleic acids, and lipids are a “nano-snapshot” of the parent cell that may be used for novel diagnostics of various diseases, including neurodegenerative disorders. Currently, diagnosis of the most common neurodegenerative movement disorder, Parkinson’s disease (PD), relies on manifestations of late-stage progression, which may furthermore associate with other neurodegenerative diseases such as progressive supranuclear palsy (PSP). Here, we profiled surface markers and other protein contents of brain-derived extracellular vesicles (bd-EVs) from PD (n= 24), PSP (n=25) and control (n=24). bdEVs displayed tetraspanins and certain microglia, astrocyte, and neuron markers, while quantitative proteomics revealed enrichment of several proteins in PD vs. control and/or PSP, including clathrin heavy chain 1 and 14-3-3 protein gamma. This characterization of EVs in the source tissue provides insights into local dynamics as well as biomarker candidates for investigation in peripheral fluids.

## INTRODUCTION

Parkinson’s disease (PD) is the most common neurodegenerative movement disorder, affecting approximately 1% of individuals over 65 years of age (de Lau & Breteler, 2006). PD is characterized by loss of pigmented neurons in the locus ceruleus and substantia nigra (Fearnley & Lees, 1991) and neuronal abnormal aggregation of alpha-synuclein (α-syn) (e.g., Lewy bodies (LBs) and Lewy neurites) (Hijaz & Volpicelli-Daley, 2020; J. Y. Li et al., 2008). Since neuronal loss in the brain is irreversible, drugs that delay or prevent neurodegeneration are more effective if prescribed at early stages of pathology. However, PD diagnosis may not occur until late, and PD can also be confused with other parkinsonian disorders, such as progressive supranuclear palsy (PSP) (Hughes et al., 2002; Joutsa et al., 2014). Therefore, reliable, specific, and easily accessed biomarkers of PD are urgently needed (Mark Frasier and Un Jung Kang, 2014).

Extracellular vesicles (EVs) already serve as biomarkers of, e.g., prostate cancer (McKiernan et al., 2020) and have elicited growing interest in the neurodegenerative disease field (Cao et al., 2019; Dutta et al., 2021; Gámez-Valero et al., 2019; Trotta et al., 2018; Wu et al., 2017). EVs are lipid bilayer-delimited nanoparticles released by various cells (Cocozza et al., 2020; van Niel et al., 2018; Yates et al., 2022), including those in the central nervous system (CNS) (Coleman & Hill, 2015; Krämer-Albers & Hill, 2016). EVs remove cellular toxins, participate in cell signaling, transfer molecular cargo, including nucleic acids and proteins, and serve as trophic factors (Maas et al., 2017; Russell et al., 2019; Yáñez-Mó et al., 2015).

In PD, peripheral biofluid EVs have been suggested as potential biomarkers, for example, for the purpose of stratifying cohorts (Alvarez-Erviti et al., 2011; Emmanouilidou et al., 2010; Stuendl et al., 2021), with the assumption that CNS EVs can escape to the periphery or that peripheral EVs otherwise reflect the state of the brain. Various PD-related proteins, such as alpha-synuclein (α-syn) (Emmanouilidou et al., 2010), LRRK2, and DJ-1 (Hong et al., 2010) are carried by EVs and, in some cases, enriched during lysosomal pathway failure (Alvarez-Erviti et al., 2011; Danzer et al., 2012). EVs have also been investigated in mitochondrial or oxidative stress and calcium dysfunction (Eitan et al., 2016; Fan et al., 2019). Although peripheral biofluid EV molecular profiles have been examined at the RNA and protein levels (Dutta et al., 2021; Jiang et al., 2021; Shi et al., 2014), how these findings compare with EVs present in the brain remains unclear. Brain-derived EVs (bdEVs) can be separated by gentle tissue digestion in a reproducible manner (Huang et al., 2020; Vella et al., 2017). Here, we separated and profiled bdEVs from PD, PSP, and control tissues. Our goals were to understand whether and how pathological conditions are reflected in EV molecular cargo and to identify molecules for capture and profiling of peripheral bdEVs.

## MATERIALS AND METHODS

*See Table 1 for manufacturers, catalog numbers, and other information*.

**Table 1.**
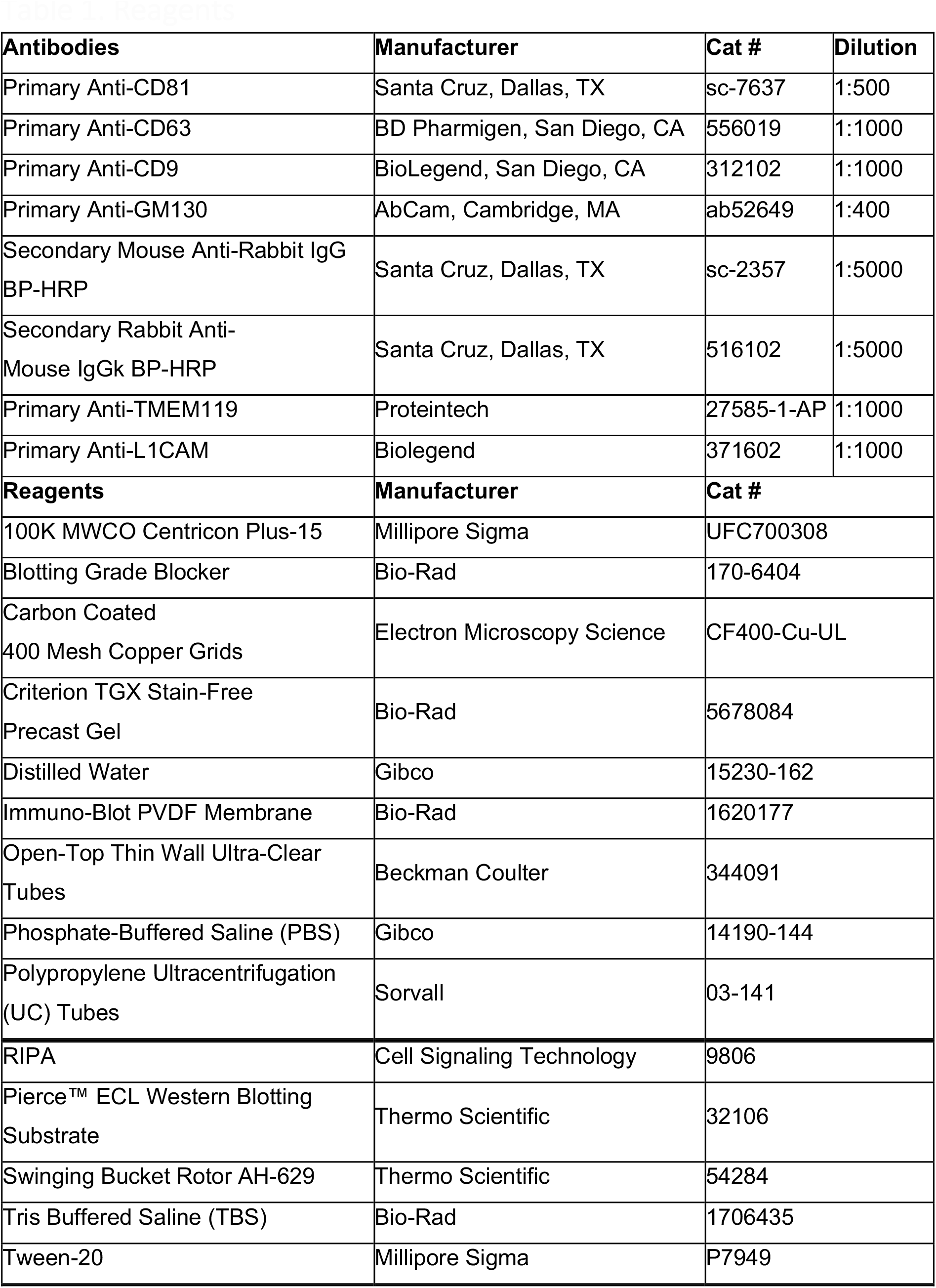
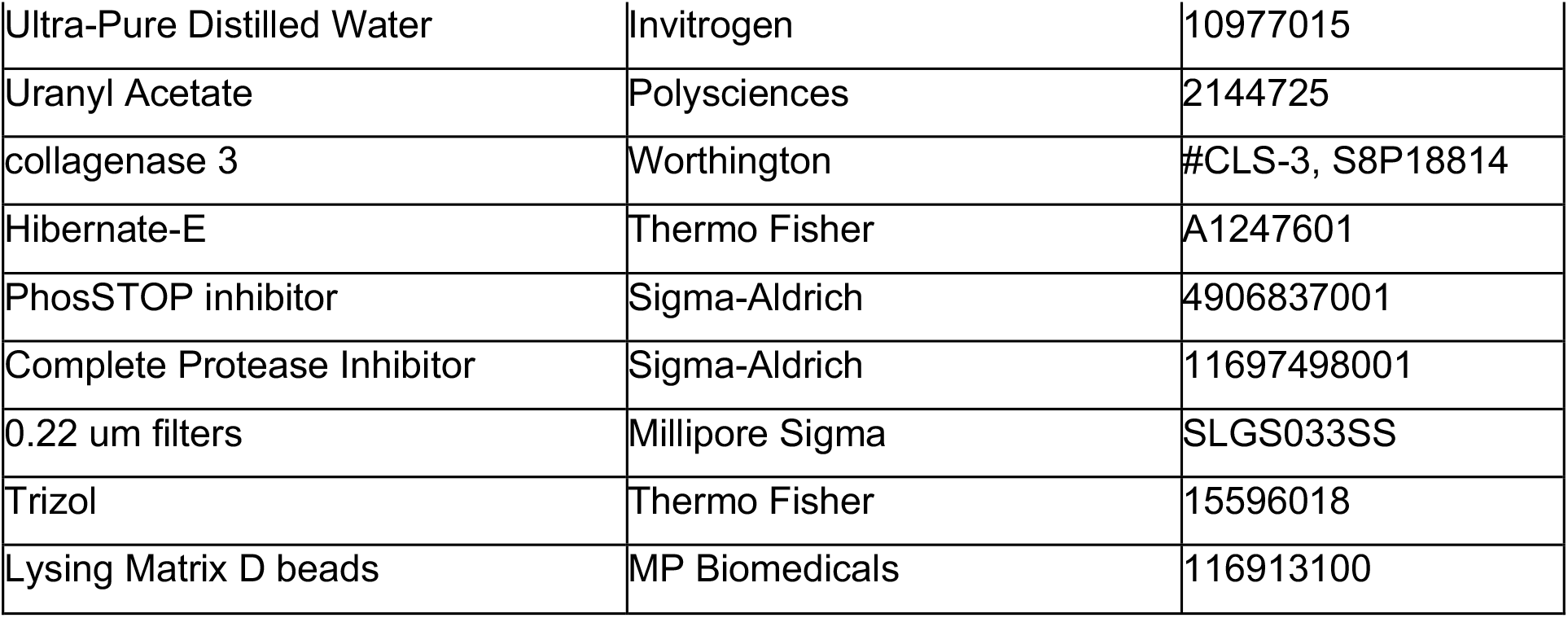
Reagents

| **Antibodies** | **Manufacturer** | **Cat #** | **Dilution** |
| --- | --- | --- | --- |
| Primary Anti-CD81 | Santa Cruz, Dallas, TX | sc-7637 | 1:500 |
| Primary Anti-CD63 | BD Pharmigen, San Diego, CA | 556019 | 1:1000 |
| Primary Anti-CD9 | BioLegend, San Diego, CA | 312102 | 1:1000 |
| Primary Anti-GM130 | AbCam, Cambridge, MA | ab52649 | 1:400 |
| Secondary Mouse Anti-Rabbit IgG BP-HRP | Santa Cruz, Dallas, TX | sc-2357 | 1:5000 |
| Secondary Rabbit Anti-Mouse IgGk BP-HRP | Santa Cruz, Dallas, TX | 516102 | 1:5000 |
| Primary Anti-TMEM119 | Proteintech | 27585-1-AP | 1:1000 |
| Primary Anti-L1CAM | Biolegend | 371602 | 1:1000 |
| **Reagents** | **Manufacturer** | **Cat #** |  |
| 100K MWCO Centricon Plus-15 | Millipore Sigma | UFC700308 |  |
| Blotting Grade Blocker | Bio-Rad | 170-6404 |  |
| Carbon Coated<br>400 Mesh Copper Grids | Electron Microscopy Science | CF400-Cu-UL |  |
| Criterion TGX Stain-Free<br>Precast Gel | Bio-Rad | 5678084 |  |
| Distilled Water | Gibco | 15230-162 |  |
| Immuno-Blot PVDF Membrane | Bio-Rad | 1620177 |  |
| Open-Top Thin Wall Ultra-Clear<br>Tubes | Beckman Coulter | 344091 |  |
| Phosphate-Buffered Saline (PBS) | Gibco | 14190-144 |  |
| Polypropylene Ultracentrifugation<br>(UC) Tubes | Sorvall | 03-141 |  |
| RIPA | Cell Signaling Technology | 9806 |  |
| Pierce™ ECL Western Blotting<br>Substrate | Thermo Scientific | 32106 |  |
| Swinging Bucket Rotor AH-629 | Thermo Scientific | 54284 |  |
| Tris Buffered Saline (TBS) | Bio-Rad | 1706435 |  |
| Tween-20 | Millipore Sigma | P7949 |  |

|  |  |  |
| --- | --- | --- |
| Ultra-Pure Distilled Water | Invitrogen | 10977015 |
| Uranyl Acetate | Polysciences | 2144725 |
| collagenase 3 | Worthington | #CLS-3, S8P18814 |
| Hibernate-E | Thermo Fisher | A1247601 |
| PhosSTOP inhibitor | Sigma-Aldrich | 4906837001 |
| Complete Protease Inhibitor | Sigma-Aldrich | 11697498001 |
| 0.22 um filters | Millipore Sigma | SLGS033SS |
| Trizol | Thermo Fisher | 15596018 |
| Lysing Matrix D beads | MP Biomedicals | 116913100 |

### Tissue collection and preparation

Human postmortem brain tissue samples were archived at −80°C by the Brain Resource Center (Department of Pathology, Johns Hopkins University School of Medicine) following brain autopsy with full neuropathological examination. For tissue dissection, neuroanatomical identification of the inferior parietal gyri was performed on coronal sections from 73 individuals (PD: n= 24; PSP: n=25; Control: n=24). The selected brain tissues were placed in a −20°C freezer for 15 minutes. Sterile 4-mm-diameter tissue punches were then used to extract samples from the inferior parietal cortex at the level of supramarginal and angular gyri. Samples were again stored at −80°C prior to bdEV separation. Several additional samples were obtained from the brain bank for the purpose of training and tool comparison. To minimize the influence of batch effects and operator bias, samples were de-identified by an individual not involved in the study and assigned to processing batches with an equal or approximately equal number of samples from each clinical group. De-identified samples were processed in these batches. Table 2 summarizes the cohort information, which was made available to investigators only after all data were obtained and archived.

**Table 2.** Subject and tissue information

|  | PD (n=24) | PSP (n=25) | Control<br>(n=24) | p-value |
| --- | --- | --- | --- | --- |
| **Age (mean)** | 79.4 +/- 7.7 | 76.1 +/- 6.8 | 77.0 +/- 9. | 0.36 |
| **Sex (male,**<br>**female)** | 9 F/ 15 M | 10 F/ 15 M | 10 F / 13 M |  |
| **PMI, mean**<br>**(hours)** | 11.9 +/- 5.8 | 11.8 +/- 8.0 | 10.7 +/- 6.0 | 0.79 |
| **Brain tissue**<br>**weight (mg)** | 416.86 +/-<br>78 | 487.00 +/-<br>127.27 | 460.05 +/-<br>96.28 | 0.052 |

### Brain-derived extracellular vesicle separation

EVs were separated from tissue as previously described (Huang et al., 2020). The frozen tissue was weighed, sliced on dry ice, and gently digested using collagenase type 3 enzyme in Hibernate-E solution for 20 min at 37°C. 1X PhosSTOP and Complete Protease Inhibitor solution (PI/PS) were added to stop the enzymatic reaction. Differential centrifugation was performed at 4°C. First, the dissociated tissue was centrifuged at 300 × g for 10 min. A small portion of each pellet (“brain homogenate with collagenase,” BHC) was stored at −80°C for later protein extraction. Supernatant was pipetted off using a 10 mL serological pipette, transferred to a fresh tube, and centrifuged at 2000 × g for 15 min. The “2K” pellets were snap-frozen and saved at - 80°C. The supernatant was further depleted of debris and large bodies through a gentle 0.22-μm filtration at a slow flow rate of 5 mL/min. Filtered material was centrifuged at 10,000 x g for 30 min using swinging-bucket rotor TH-641 (Thermo Scientific, k-factor 114, acceleration and deceleration settings of 9). Pellets (“10K”) were resuspended in 150 μl PBS containing 1X PI/PS by pipetting up and down ten times, vortexing 30 seconds, and incubating on ice 20 minutes, followed by a repeat of the pipetting and vortexing steps. Thusly resuspended 10K bdEVs were aliquoted and stored at −80°C. Supernatants were transferred into 100 kilodaltons (kDa) molecular weight cut-off (MWCO) protein concentrators (Thermo Fisher, 88524), and volumes were reduced from 5 mL to 0.5 mL. Retentate was then applied to the top of qEV Original size exclusion chromatography (SEC) columns (Izon Science, Cambridge, MA) that were pre-rinsed with 15 ml PBS. 0.5 mL fractions were collected by elution with PBS, using Izon automated fraction collectors (AFCs; Izon Science, Cambridge, MA). Fractions 1-6 (≈ 2.8 mL) were considered the void volume; fractions 7-10 were pooled as EV-enriched fractions; and fractions 11-14 were pooled as protein-enriched fractions. EV-enriched fractions were transferred to polypropylene UC tubes and centrifuged at 100,000 × g for 70 min, using the TH-641 swinging-bucket rotor as described above. Supernatant was poured off, and UC tubes were stood upright and inverted on a piece of tissue to remove residual buffer. Pellets (“100K”) were resuspended following the same procedure described above, but by adding 120 μl PBS (1X PI/PS) to each tube. Aliquots were stored at − 80°C.

### Nano-flow cytometry measurement (NFCM)

Particle concentration and size profiles were assessed by nano-flow (Flow NanoAnalyzer, NanoFCM, Nottingham, England) using side scatter, as described previously (Arab et al., 2021; Huang et al., 2020). Instrument calibrations for concentration and size distribution were done using 200-nm polystyrene beads and a silica nanosphere mixture (68-155 S16 M-EXO), respectively. After calibration, settings remained unchanged during sample evaluation. For each sample, 2 μl of bdEV resuspension was used. Serial dilutions were performed as needed to fall within the manufacturer’s recommended particle count range (Arab et al., 2021), and events were recorded for 1 min. Flow rate and side-scatter intensity were converted into particle concentration and size distribution using calibration curves.

### Transmission electron microscopy (TEM)

12 of 73 samples were randomly chosen and imaged by TEM as previously described (Arab et al., 2021). Briefly, 10 μl of each sample was freshly thawed and adsorbed to glow-discharged carbon-coated 400 mesh copper grids by flotation for 2 min. Three consecutive drops of 1× Tris-buffered saline were prepared on Parafilm. Grids were washed by moving from one drop to another, with a flotation time of 1 min on each drop. The rinsed grids were then negatively stained with 1% uranyl acetate (UAT) with tylose (1% UAT in deionized water (dIH2O), double-filtered through a 0.22-μm filter). Grids were blotted, then excess UAT was aspirated, leaving a thin layer of stain. Grids were imaged on a Hitachi 7600 TEM operating at 80 kV with an XR80 charge-coupled device (8 megapixels, AMT Imaging, Woburn, MA, USA).

### Western blot (WB)

BH, BHC, 2K, 10K, and EV and protein SEC fractions were lysed in 1× radioimmunoprecipitation assay buffer (RIPA) supplemented with protease inhibitor cocktail. A total of 20 μL of lysates were resolved using a 4% to 15% Criterion TGX Stain-Free Precast gel, then transferred onto an Immuno-Blot PVDF membrane. Antibodies to CD81, CD63, and CD9 were used to detect EV membrane markers, and anti-GM130 antibody was used to detect Golgi intracellular contamination. Antibodies were diluted in PBS-T containing 5% blotting-grade blocker (Bio-Rad, #1706404). Membranes were incubated overnight (≈16 h). After several washes in PBS-T, rabbit anti-mouse IgGk BP-HRP and mouse anti-rabbit IgGk BP-HRP secondary antibodies were diluted in blocking buffer, and membranes were incubated for 1 h at room temperature (RT). Pierce™ ECL Western Blotting Substrate (Thermo Fisher, 32106) was applied, and blots were visualized using a Thermo Fisher iBright 1500 imaging system.

### Single-particle interferometric reflectance imaging sensor (SP-IRIS)

EVs were phenotyped with EV-TETRA-C ExoView Tetraspanin kits and an ExoView TMR100 scanner (NanoView Biosciences, Boston, MA) according to the manufacturer’s instructions and as described previously (Arab et al., 2021). Concentrations as measured by NFCM, were adjusted such that around 1e9 particles from each sample were mixed 1:1 with ExoView incubation buffer (IB). 35μl of this mixture was placed onto the chip and incubated at RT for 16h (no shaking). Chips were washed with IB and incubated 1h at RT with a fluorescently-labeled antibody cocktail of anti-human CD81 (JS-81, CF555), CD63 (H5C6, CF647), and CD9 (HI9a, CF488A) at dilutions of 1:1200 (v:v) in a 1:1 (v:v) mixture of IB and blocking buffer. All chips were scanned and imaged with the ExoView scanner using both SP-IRIS Single Particle Interferometric Reflectance Imaging Sensor and fluorescence detection. Data were analyzed using NanoViewer 2.8.10 Software.

### Multiplexed ELISA

Prototype S-PLEX® ultrasensitive assays (Meso Scale Diagnostics, Rockville, MD) were used for intact EVs. Five multiplexed assay panels were assembled in this fashion. According to the manufacturer’s recommendations, samples were diluted up to 30-fold in “diluent 52,” added to the plates, and incubated at RT with continuous shaking. Panel 1, comprising antibodies targeting relatively abundant surface markers, was incubated for 1 hour, while the remaining panels, targeting lower-abundance markers, were incubated for 4 hours to improve sensitivity. EVs captured by each antibody were detected using MSD’s S-PLEX® ultrasensitive assay methods with a cocktail of detection antibodies targeting CD63, CD81, and CD9. Assay plates were read with MSD GOLD™ Read buffer B on an MSD® SECTOR instrument. bdEVs from the 73 subjects, as well as additional positive and negative controls, were assayed with each panel.

### Relative quantification proteomics

#### Sample preparation, protein digest, peptide TMT labeling

Proteins were extracted from 20 μl of resuspended bdEVs separated from the inferior parietal region of Parkinson’s (n=5), Progressive Supranuclear Palsy (n=5), and matched controls samples (n=5) using RIPA buffer (Cell Signaling Technology). Disulfide bonds were reduced using 50 mM dithiothreitol (Sigma, D0632) and cysteine residues were alkylated with iodoacetamide (Sigma, I1149) as previously described (Arab et al., 2019). Resulting protein lysates were precipitated with trichloroacetic acid (TCA) (Sigma, T0699). Briefly, samples were supplemented with 10% of TCA in acetone solution in a ratio of (1:8, V/V). After precipitation, pellets were resuspended in 50 mM ammonium bicarbonate containing 10% acetonitrile, sonicated for 10 min, and pH was adjusted to 7.8–8.0 using Tris Hydrochloride, pH 8.0. Proteins were digested with 12 ng/mL Trypsin/Lys-C (Thermo Fisher Scientific # A41007) overnight at 37 °C. Digested peptides were labeled with 16-plex TMT isobaric mass tag reagents (Thermo Fisher Scientific) according to the manufacturer’s instructions. Labeled peptides were loaded onto a Pierce detergent removal column (Thermo Fisher # 87779) and eluted in 300 μL of 8% Triethylammonium bicarbonate, 0.1% Trifluoroacetic acid, and 92% water. Prior to mass spectrometry analysis, peptides were desalted on u-HLB Oasis plates (Waters), eluted in a buffer containing 60% acetonitrile and 0.1%TFA, and dried by vacuum centrifugation.

#### Liquid chromatography separation and tandem mass spectrometry (LC-MS/MS)

Desalted peptides were reconstituted in 40 μL solution buffer containing 2% ACN and 0.1% FA andanalyzed by nano-LC tandem mass spectrometry (LC-MS/MS) using a Orbitrap-Lumos-Fusion mass spectrometer (Thermo Fisher Scientific) interfaced with a EasyLC1200 series equipped with ProntoSIL-120-5-C18 H BISCHOFF reverse phase column (75 μm × 150 mm, 3 μm). Peptides were separated using a 2%–90% acetonitrile gradient in 0.1% FA over 78 min at a flow rate of 300 nL/min. Eluting peptides were electrosprayed at 2.4 kV into an Orbitrap-Lumos-Fusion mass spectrometer through a 1 μm emitter tip (New Objective). Survey scans (MS1) of peptides between 375 and 1600 m/z were acquired at 120,000 FWHM resolution (at 200 m/z) and 4e5 automatic gain control (AGC). By data-dependent acquisition, the 15 most intense ions were individually isolated with 0.7 m/z window and fragmented (MS/MS) using a 38 higher-energy collisional dissociation (HCD) activation energy and 15 sec dynamic exclusion. Fragment ions were analyzed at a resolution of 60,000 FWHM and 5e4 AGC.

#### Raw data processing and analysis

All the tandem MS/MS spectra (signal/noise >2) were processed by Proteome Discoverer (v2.4 Thermo Fisher Scientific). MS/MS spectra were searched with Mascot v.2.6.2 (Matrix Science, London, UK), and proteins were identified by searching against the 2021_204_H_sapiens database. Trypsin specificity was used for the digestion, with up to two missed cleavages. Methionine oxidation and deamidation were selected as the variable modifications. Carbamidomethylation of cysteines and TMT-label modifications of N-termini and lysine were set as fixed modifications. Based on a concatenated decoy database search, Proteome Discoverer-Percolator was used to validate identified peptides with a confidence threshold of 1%. Proteome Discoverer used TMT reporter ions from peptide-matched spectrum (PSM) of unique Rank 1 unmodified peptides with reporter signal/noise >5 and an isolation interference <25 to calculate and normalize protein ratios based on the normalized median ratio of all spectra of tagged peptides from the same protein (Herbrich et al., 2013). Gene ontology terms were determined using PANTHER V17.0 software

### Statistical analysis and data and methods availability

Differences in total EV particle concentrations and in surface markers expression were assessed by Kruskal-Wallis ANOVA test in GraphPad Prism 9 (GraphPad Software Inc., La Jolla, CA). Results were considered significant if p-value < 0.05. For labeled proteins, significant differential abundance was defined as Log 2-fold-change = 0.32 and p-value <0.05. p-values and adjusted p-values (Benjamini-Hochberg method) were calculated using a non-nested ANOVA test per Proteome Discoverer 2.4 recommendations.

All relevant experimental details were submitted to the EV-TRACK knowledgebase (EV-TRACK ID: EV220312) (van Deun et al., 2017). Proteomic data are available upon request.

## RESULTS

### bdEV separation and characterization

bdEVs were separated from brain tissue of a selected cohort of Parkinson’s, PSP, and control donors (PD: n= 24; PSP: n=25; Control: n=24) in compliance with the international consensus guidelines for studies of EVs (MISEV2018) (Théry et al., 2019). Characterization of specific markers included Western blot and SP-IRIS for several samples with greater availability, and, for more precious cohort samples, proteomics and multiplexed ELISA. For selected samples, Western blot (WB) showed low levels of CD63 but abundant CD9 and CD81; Golgi marker GM130 was relatively depleted (Fig. 2a). Similarly, SP-IRIS confirmed that bdEVs were positive for the tetraspanins CD81, CD63, and CD9, along with cytosolic EV marker Syntenin (Fig 2b). Fluorescence signals for CD63 and Syntenin were similar across tetraspanin capture spots, whereas fluorescence for CD9 and CD81 varied based with capture antibody (Sup Fig. 1 c-g).

**Figure 1.**
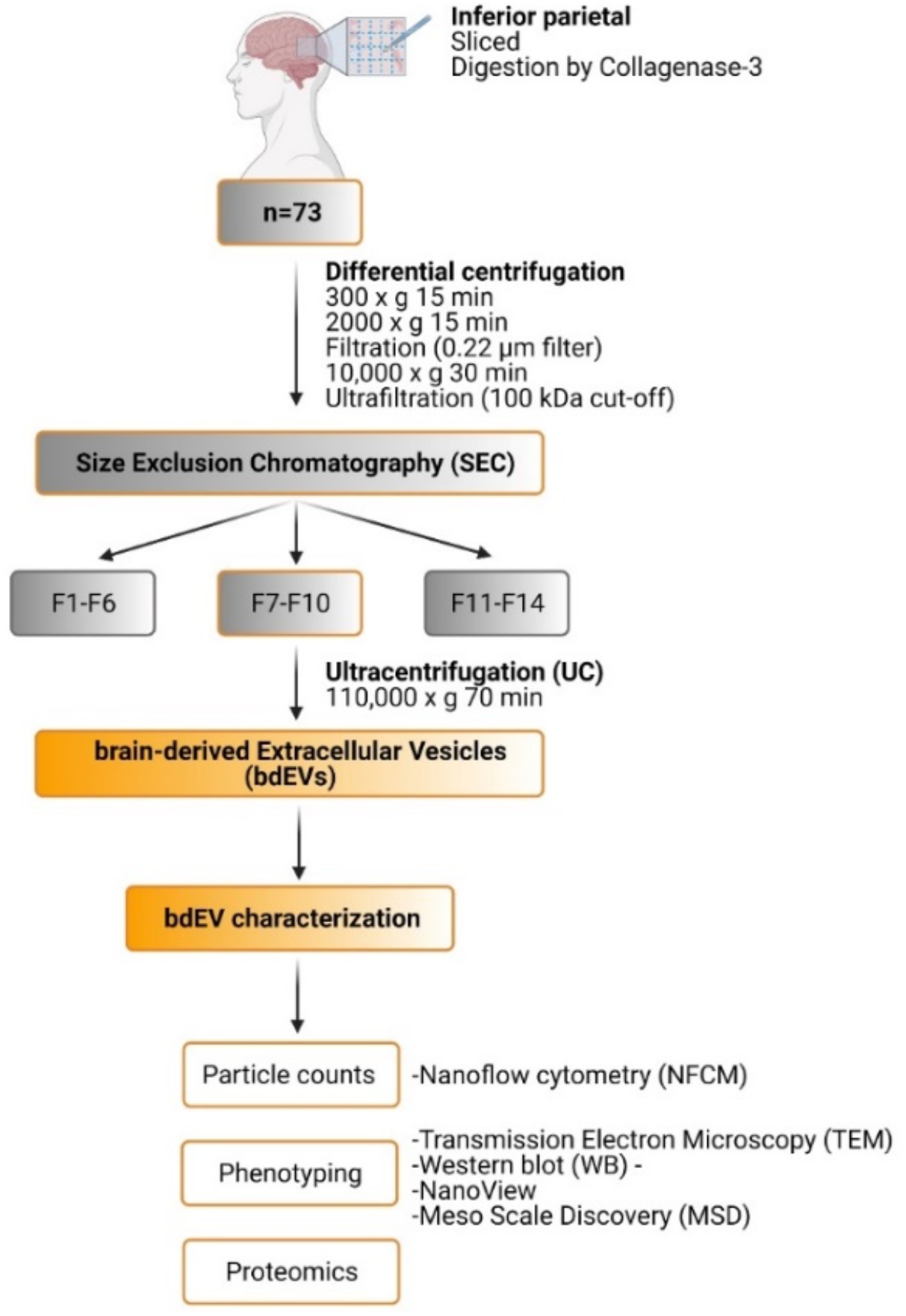
bdEV separation and characterization workflow. Inferior parietal tissues from 73 human subjects were sliced into small pieces and gently digested. The resulting bdEVs were assessed for particle counts and size distribution using nanoFCM. Further characterizations were done with electron microscopy, Western blot, Single Particle Interferometric Reflectance Imaging Sensor (SP-IRIS), multiplexed ELISA (Meso Scale Discovery), and proteomics. Figure created with BioRender.

**Figure 2.**
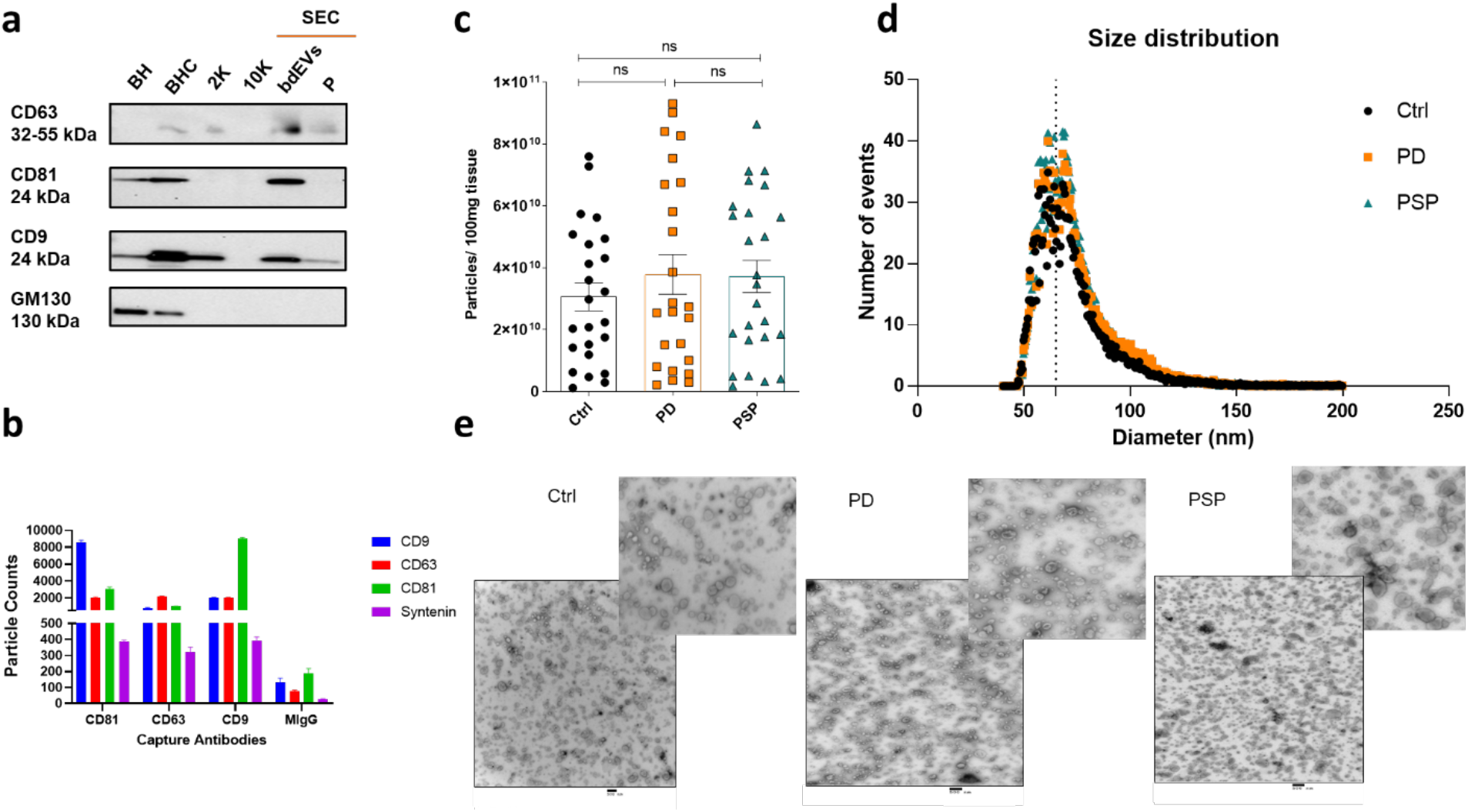
bdEV characterization. (a) Western blots of EV surface markers CD63, CD81, and CD9, and Golgi protein GM130 (BH = brain homogenate, BHC = brain homogenate plus collagenase, 2K and 10K = centrifugation pellets, SEC = size exclusion chromatography, with EV fractions and protein (P) fractions. (b). SP-IRIS (fluorescence) detection of tetraspanins (CD9, CD63, CD81) and cytosolic syntenin. Results are presented as mean +/- SD of three capture spots as indicated (CD81, CD63, CD9, and IgG negative control). (c) Particle concentration of human bdEVs by NFCM for control (black), PD (orange), and PSP (green) normalized by tissue input. Data are presented as mean +/- SD. ns: no significant difference (P>0.05), Kruskal-Wallis ANOVA test. (d) Diameter distribution profile (NFCM). (e) Transmission electron micrographs (TEM, scale bar = 500 nm).

For all cohort samples, particle counts and size distribution per 100 mg of tissue were measured using NFCM. There were no significant differences in particle counts (Fig. 2c), size distribution profiles (Fig. 2d), or mean size (Fig. supp 1b) between PD, PSP, and control groups. Diameter distribution profiles for each group displayed the same pattern, with a peak around 65 nm. TEM images of randomly selected samples across control, PD, and PSP groups showed particles with expected EV morphology and size (Fig. 2c). Consistent with NFCM, there were no differences in size and yield by group (Fig. 2c, supp Fig. 1a). Additional characterization by mass spectrometry and ELISA surface phenotyping is presented below.

### bdEV, tetraspanin, and cell marker phenotyping suggest disease-associated differences

To assess the relative contribution of each brain cell type to total bdEVs, multiplexed ELISAs were used to assay 36 proteins, including markers of one or more specific cell types: microglia, neurons, astrocytes, and endothelial cells (Fig. 3a).

**Figure 3.**
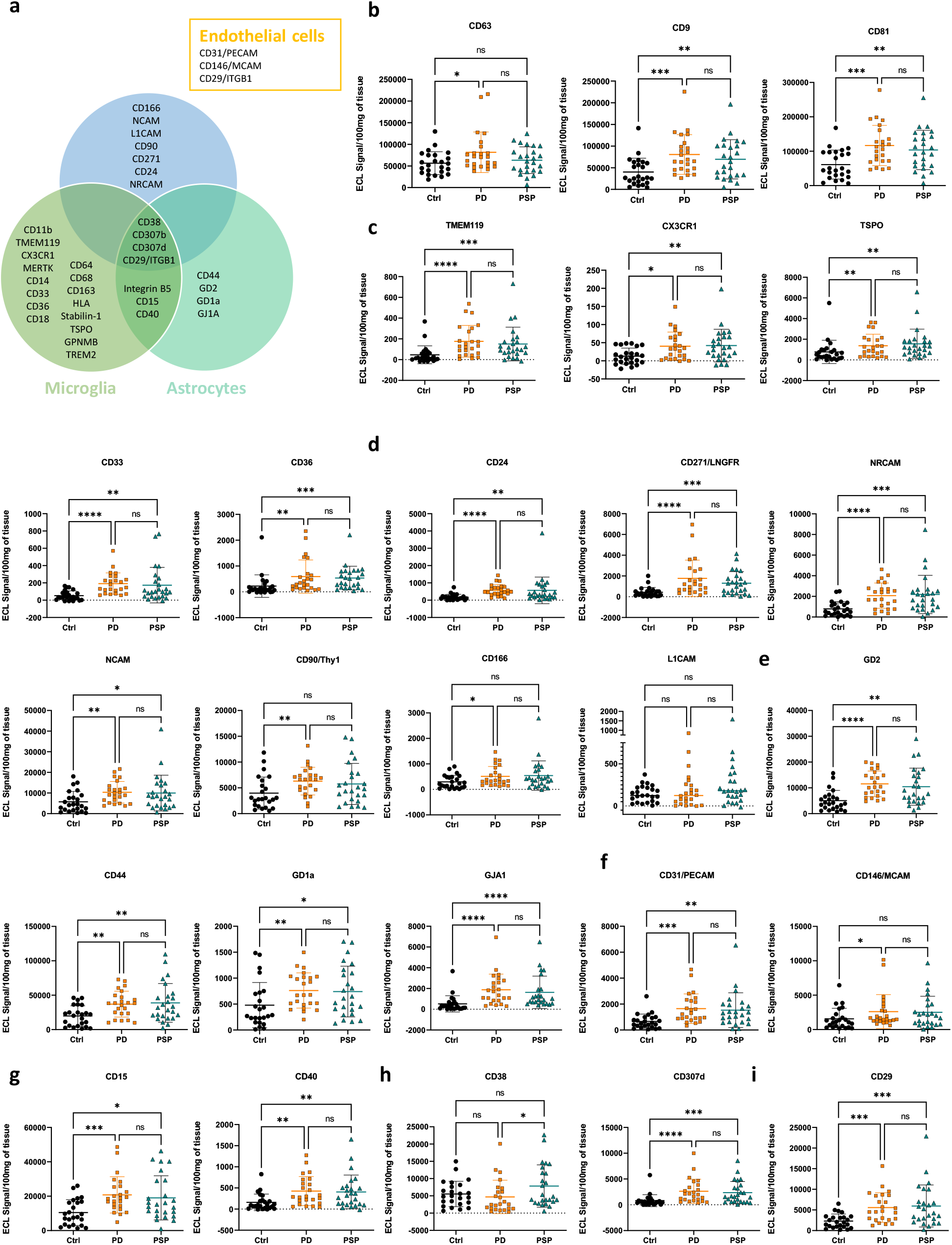
bdEV phenotyping by multiplexed ELISA. (a) Distribution of markers by cell type: microglia, neurons, astrocytes, and endothelial cells. Comparison of control (Ctrl), PD, and PSP groups for (b) general markers of EVs and markers of cell of origin: (c) microglia, (d) neurons, (e) astrocytes, (f) endothelial cells; and (g, h, i) markers of multiple populations as indicated. Data were normalized per 100 mg of tissue input and reported as mean +/- SD. Ctrl (n=24); PD (n=24); PSP (n=25). * = p < 0.05, ** = p < 0.01, *** = p < 0.001, **** = p < 0.0001, and ns = non-significant (Kruskal-Wallis ANOVA).

### Tetraspanins

CD63 signal was significantly greater in PD compared with the control group. CD81 and CD9 signals were significantly higher for PD and PSP than controls (Fig. 3b), but there were no differences between the disease groups. Several cellular origin surface markers were increased in bdEVs from PD patients compared with control (Fig. 3).

### Microglial markers

Transmembrane protein 119 (TMEM119), C-X3-C motif chemokine receptor 1 (CX3CR1), translocator protein (TSPO), CD33, and CD36 were significantly more abundant in PD and PSP compared with control, but the difference between PD and PSP was not statistically significant (Fig. 3c). Stabillin-1 differentiated PSP from control and glycoprotein nonmetastatic melanoma protein B (GPNMB) was different between PD from control (Supp Fig. 3a; note that some values are negative due to background subtraction). CD64, major histocompatibility complex class II (HLA-DR/DP/DQ), CD163, CD68, and CD32b showed no significant difference between groups (Supp Fig. 2a). Triggering receptor expressed on myeloid cells 2 (TREM2), CD11b, CD18, and MER proto-oncogene, tyrosine kinase (MERTK) were below the limit of detection (data not shown) ITGB5.

### Neuronal markers

CD24, nerve growth factor receptor (CD271), and neuronal cell adhesion molecules NRCAM and NCAM were significantly different for disease groups compared with controls, but PD did not differ from PSP. Thy-1 cell surface antigen (CD90/Thy1) and CD166 differed between PD and control but not between PSP and control. Only low levels of signal for L1 cell adhesion molecule (L1CAM) were detected, with no significant differences between groups (Fig. 3d).

### Astrocytic markers

Ganglioside G2 (GD2) and CD44 were detected abundantly compared with ganglioside GD1a (GD1a) and gap junction alpha-1 protein (GJA1, Fig. 3e). These markers show significant differences between the pathological groups compared with control.

### Endothelial markers

Platelet and endothelial cell adhesion molecule 1 (CD31) differed in the pathological groups compared with the control group, whereas melanoma cell adhesion molecule (CD146) differentiated PD, but not PSP, from control (Fig. 3f).

### Non-tetraspanin proteins found in multiple cell types

Fucosyl transferase 4 (CD15) and CD40, found on both microglia and astrocytes, distinguished both disease groups from control (Fig. 3g), whereas integrin subunit beta 5 (ITGB5) showed no significant differences between tested groups (Supp Fig. 2b). CD38 (cluster of differentiation 38), also known as cyclic ADP ribose hydrolase, and Fc receptor-like 4 (CD307d), both of which are found on microglia, astrocytes, and neurons, differentiated PD from PSP and pathological groups from the control group, respectively (Fig. 3h). CD307d showed values close to background for the majority of tested samples (Supp Fig. 2 c). Integrin subunit beta 1 (CD29), a marker detected in all cells, distinguished both disease groups from control (Fig. 3 i).

### Differentially expressed proteins in brain-derived EVs as revealed by quantitative proteomics

Relative quantitative mass spectrometry was performed for bdEVs from five individuals in each group (PD, PSP, and control). A total of 1369 proteins were detected in at least one sample. Amongst these proteins, the gene ontology (GO) term “transport” was the most represented (Supp Fig. 3a). 306 master proteins were successfully tagged for quantitative comparison. After removing duplicates, per calculated ratios PD vs. control, PD vs. PSP, and PSP vs. control, 26 proteins were identified as significantly differentially abundant (Fig. 4a, Table 3). In this list, there are 17 proteins less abundant by Log 2-fold-change (Log_2_ fc) less than or equal to - 0.32 and p-value <0.05, and 7 proteins more abundant by Log_2_ fc greater than or equal to 0.32 and p-value <0.05 (Supp Fig. 3b-d). Ankyrin-1 (ANK1) and stomatin (STOM) were more abundant in PSP vs. control and less abundant in PD vs. PSP.

**Table 3.** Differentially abundant proteins

| Gene Symbol | PD/C |  | PSP/C |  | PD / PSP |  |
| --- | --- | --- | --- | --- | --- | --- |
|  | Abundance Ratio | Abundance Ratio p-value | Abundance Ratio | Abundance Ratio p-value | Abundance Ratio | Abundance Ratio p-value |
| HAPLN2 | 1.89 | 0.004 | 0.88 | 0.592 | 2.16 | 0.001 |
| SLC8A2 | 0.6 | 0.009 | 1.13 | 0.739 | 0.53 | 0.034 |
| GNG3 | 0.37 | 0.009 | 0.71 | 0.582 | 0.53 | 0.055 |
| ACTR2 | 0.57 | 0.012 | 1.01 | 0.865 | 0.57 | 0.021 |
| RPS9 | 0.31 | 0.016 | 0.83 | 0.992 | 0.37 | 0.028 |
| RALA | 0.69 | 0.024 | 0.66 | 0.041 | 1.04 | 0.94 |
| GNB2 | 0.66 | 0.025 | 0.81 | 0.471 | 0.81 | 0.194 |
| ATP2B1 | 0.68 | 0.026 | 1.19 | 0.774 | 0.57 | 0.008 |
| BCAN | 0.65 | 0.027 | 1.05 | 0.994 | 0.61 | 0.022 |
| RPS18 | 0.55 | 0.027 | 1.02 | 0.992 | 0.54 | 0.033 |
| GFAP | 0.38 | 0.028 | 0.87 | 0.984 | 0.43 | 0.038 |
| SLC4A1 | 2.25 | 0.033 | 2.56 | 0.012 | 0.88 | 0.843 |
| RAP1B | 1.58 | 0.035 | 1.18 | 0.996 | 1.34 | 0.04 |
| RPL15 | 0.26 | 0.036 | 0.73 | 0.995 | 0.35 | 0.043 |
| YWHAG | 0.49 | 0.043 | 1.28 | 0.908 | 0.38 | 0.02 |
| ATP2B2 | 0.6 | 0.048 | 0.72 | 0.191 | 0.82 | 0.696 |
| ARF5 | 0.65 | 0.069 | 1.08 | 0.971 | 0.6 | 0.046 |
| EPB42 | 1.9 | 0.088 | 2.92 | 0.038 | 0.65 | 0.785 |
| CAPZA1 | 0.82 | 0.105 | 1.31 | 0.127 | 0.63 | 0.002 |
| ANK1 | 1.93 | 0.11 | 4.72 | 0.001 | 0.41 | 0.032 |
| SLC1A3 | 1.36 | 0.116 | 1.7 | 0.027 | 0.8 | 0.689 |
| SLC2A1 | 1.94 | 0.134 | 2.37 | 0.026 | 0.82 | 0.624 |
| SV2A | 0.54 | 0.139 | 1.17 | 0.901 | 0.46 | 0.052 |
| STOM | 1.46 | 0.145 | 2.68 | 0.001 | 0.54 | 0.042 |
| CLTC | 0.75 | 0.169 | 1.22 | 0.222 | 0.62 | 0.008 |
| CTSD | 1.05 | 0.501 | 0.87 | 0.308 | 1.2 | 0.048 |

Light blue: considered significantly less abundant
Light red: considered significantly more abundant

**Figure 4.**
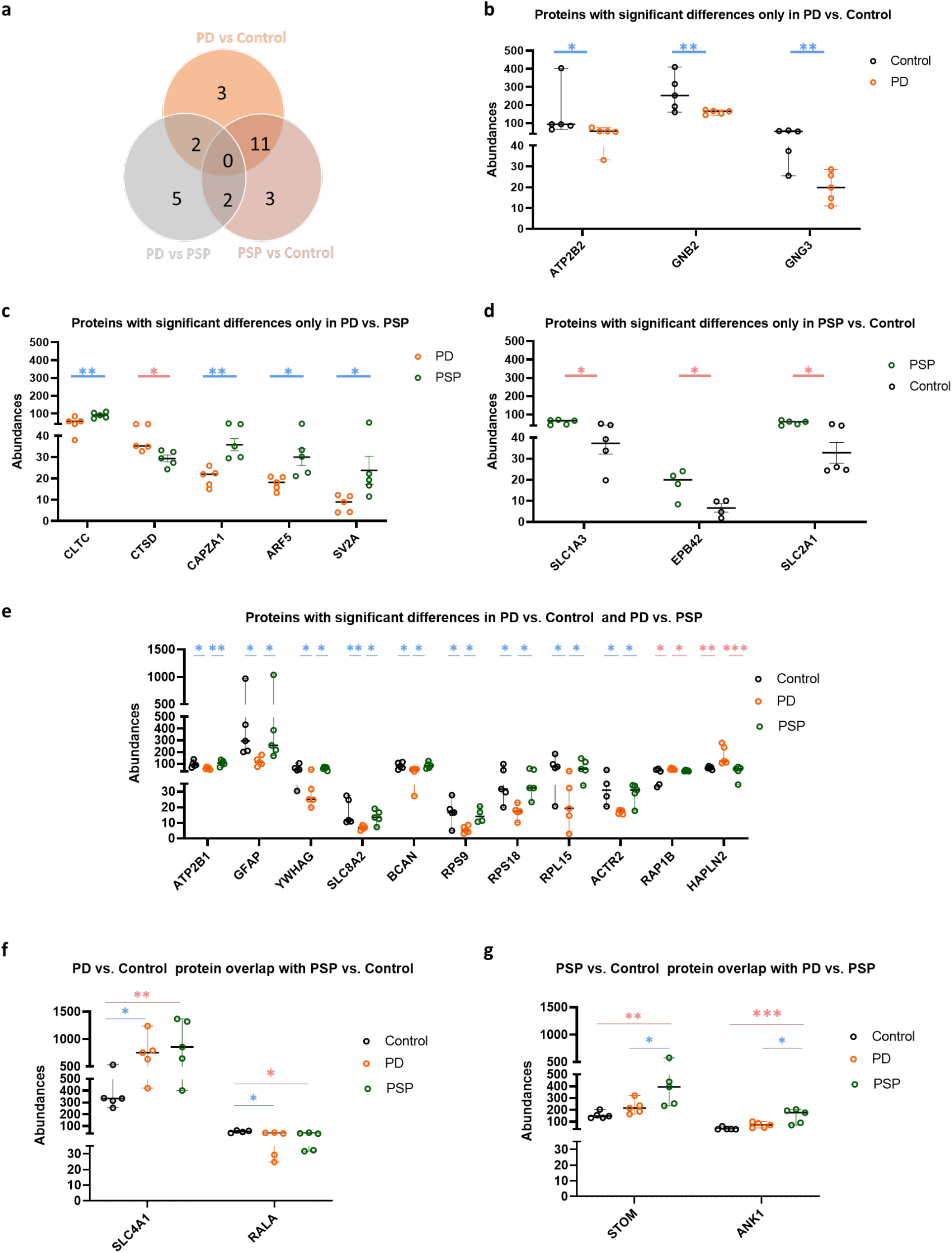
Differential proteomic cargo in bdEVs. (a) Venn diagram of the number of proteins identified as differentially abundant in multi-group comparison (p < 0.05 and log2 fold change > 0.32). Protein abundance was normalized by total identified peptides. (b-g) Normalized abundance in multi-group comparisons as indicated. Data are presented as mean +/- SD in control, PD, and PSP. * = p < 0.05, ** = p < 0.01, *** = p < 0.001 following ANOVA. Light blue bars: considered significantly less abundant. Light red bars: considered significantly more abundant.

Plasma membrane calcium-transporting ATPase 2 (ATP2B2), guanine nucleotide-binding protein G subunit beta-2 (GNB2), and guanine nucleotide-binding protein G subunit gamma-3 (GNG3) differed only between PD vs. control, with lower abundance in PD (Fig. 4b). Five proteins differed only between PD and PSP (Fig. 4c). Clathrin heavy chain 1 (CLTC), F-actin-capping protein subunit alpha-1 (CAPZA1), ADP-ribosylation factor 5 (ARF5), synaptic vesicle glycoprotein 2A (SV2A) were less abundant in PD, whereas cathepsin D (CTSD) was more abundant. Comparison between PSP and control returned three unique proteins more abundant in PSP; namely excitatory amino acid transporter 1 (SLC1A3), erythrocyte membrane protein band 4.2 (EPB42), and solute carrier family 2 (SLC2A1) (Fig. 4d).

Eleven proteins overlapped between PD vs. control and PD vs. PSP (Fig. 4e). Calcium-transporting ATPase 1 (ATP2B1), glial fibrillary acidic protein (GFAP), 14-3-3 GAMMA (YWHAG), sodium/calcium exchanger 2 (SLC8A2), brevican core protein (BCAN), ribosomal protein S9 (RPS9), ribosomal protein S18 (RPS18), ribosomal protein L15 (RPL15), actin-related protein 2 (ACTR2) were less abundant in PD, whereas Ras-related protein Rap-1b (Rap1b) and hyaluronan and proteoglycan link protein 2 (HAPLN2) were more abundant in PD. Two proteins overlapped between PD vs. control and PSP vs. control (Fig. 4f), and two proteins overlapped between PD vs. PSP and PSP vs. control (Fig. 4g). Although our mass spectrometry protocol was not directed towards membrane proteins, several membrane markers measured by ELISA were also detected by mass spectrometry (Table 4), including astrocyte marker CD44 and neuronal marker NCAM1. EV marker CD81 was detected but not TMT labeled.

**Table 4:**
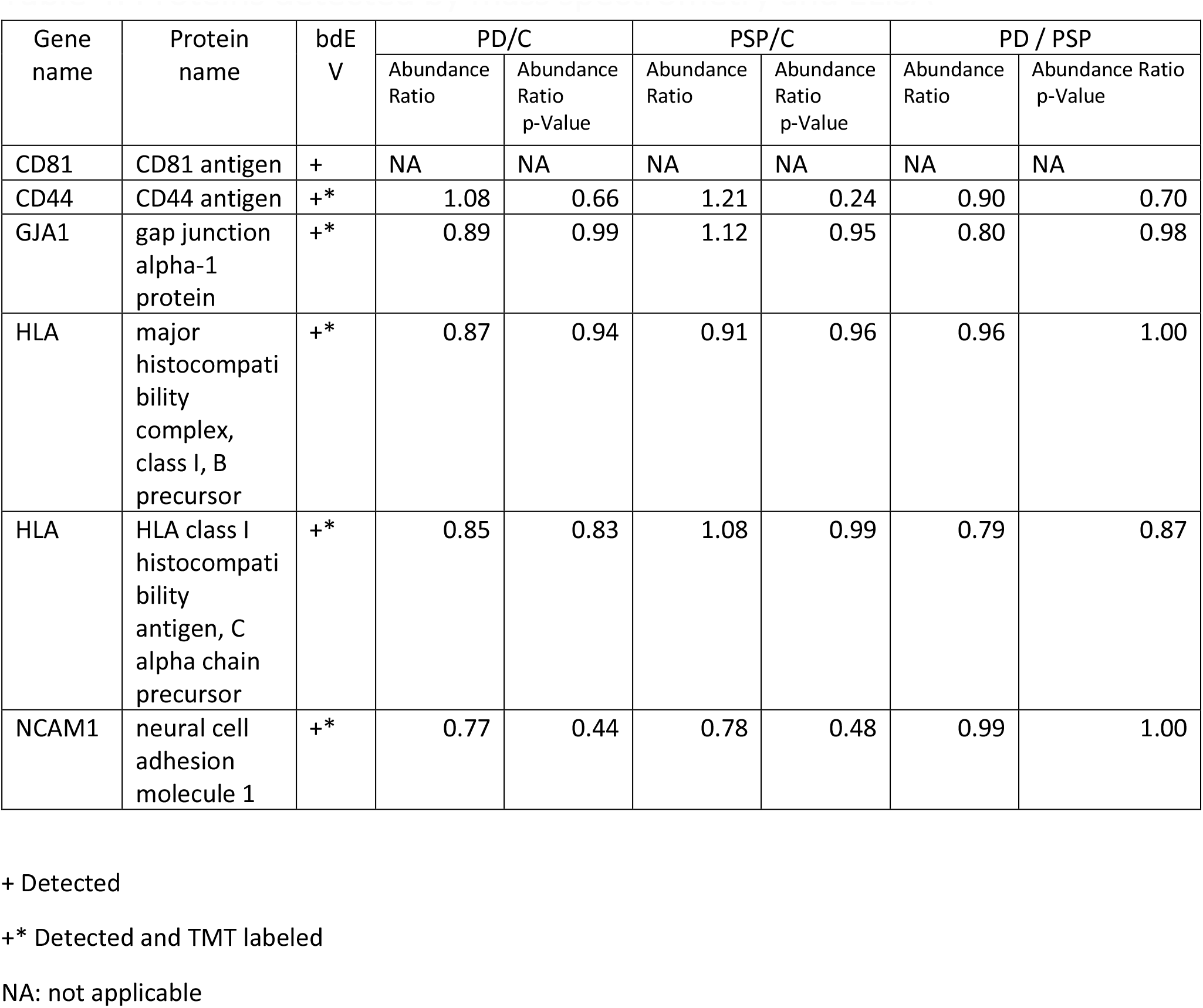
Proteins detected by mass spectrometry and ELISA

### Presence of classical EV markers and Parkinson’s disease-related proteins in brain-derived extracellular vesicles from Parkinson’s subjects

Next, we compared our proteomic data set to the ExoCarta “Top 100” list of putative EV-associated proteins. 47 proteins were present in our samples, with 37 successfully labeled with TMT tags and 10 not labeled (Table 5). Some have already been listed as distinguishing bdEV from the tested groups, e.g., YWHAG, STOM, GNB2, RAP1B, and CLTC. More EV markers were identified but without statistically significant differences between tested groups, e.g., CD81, flotillin-1 (Flot-1), RAB proteins (RAB1A, RAB5c), annexins (ANXA1, ANXA11, ANXA2, ANXA4, ANXA5, ANXA6), enolase-1 (Eno1), and 14-3-3 proteins (YWHAE, YWHAH, YWHAQ, YWHAZ). In addition, flotillin-2 (Flot-2) and syntenin-1 EV markers are present in our data set but not in the top-100 ExoCarta list.

**Table 5:**
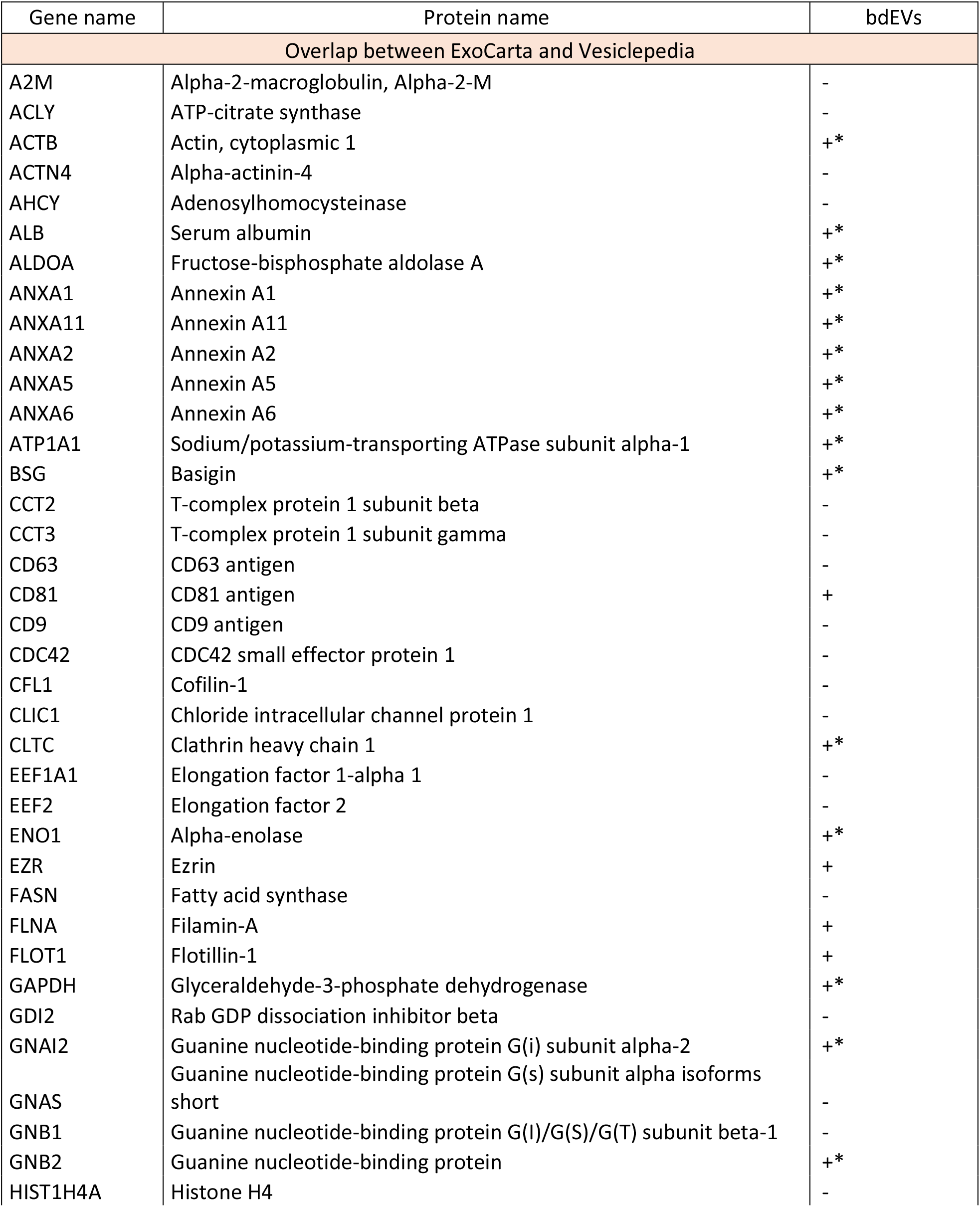
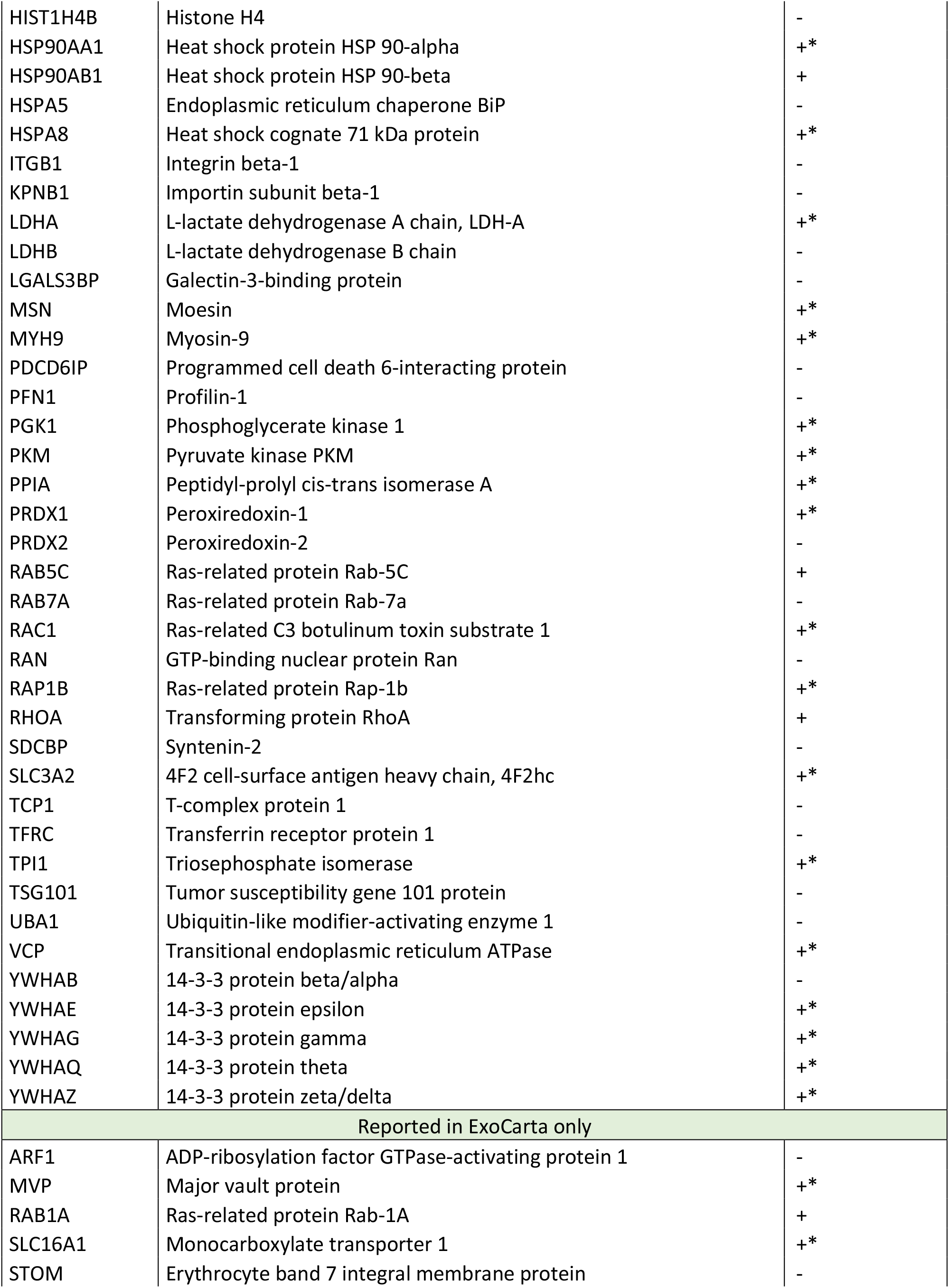
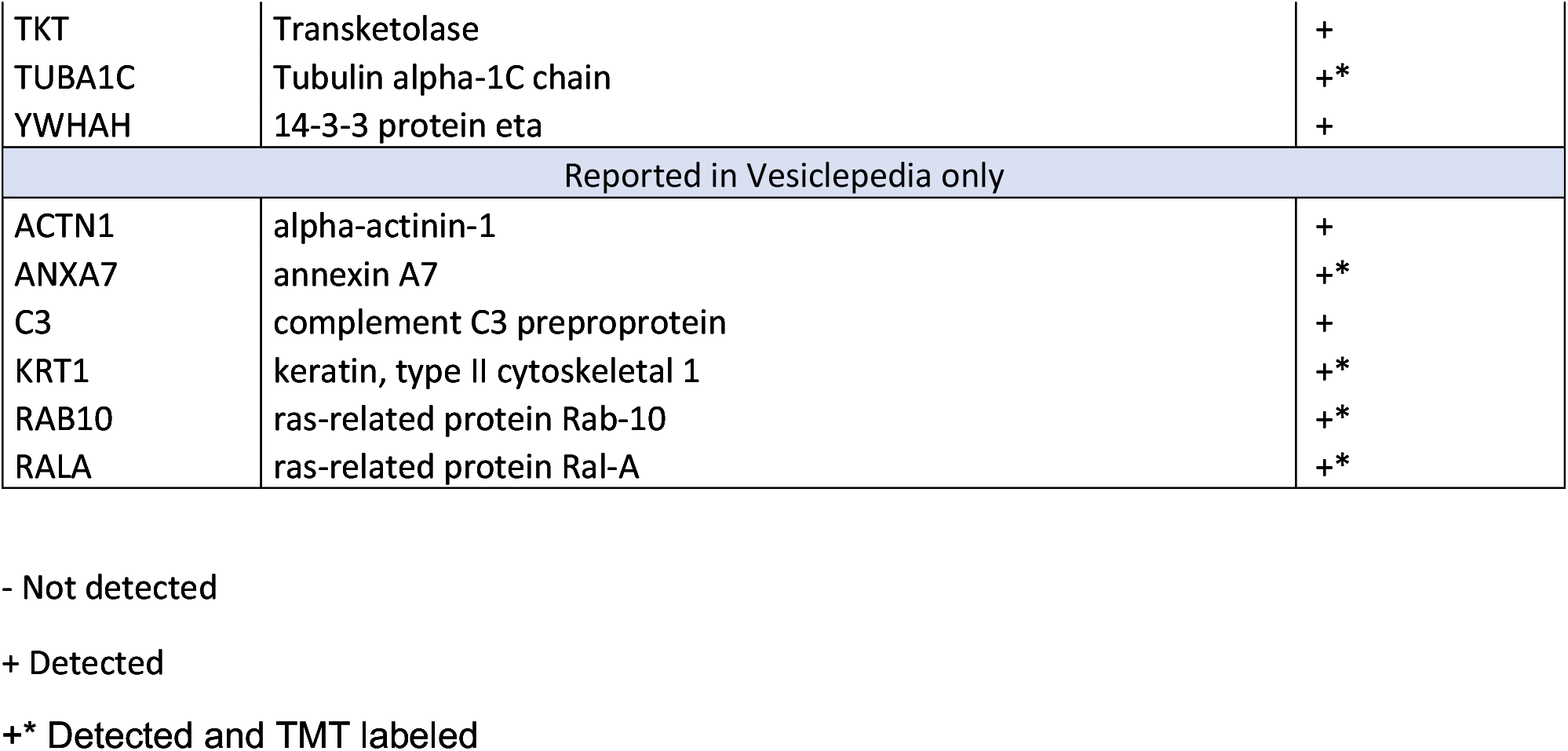
Proteins identified in bdEVs and in the “Top 100” lists of ExoCarta and/or Vesiclepedia

We followed a similar approach and compared a list of 1952 known PD genes downloaded from DisGeNET with our data set (Supp Fig. 4a). We found 80 to overlap with identified proteins, including 58 that were successfully labeled. Interestingly, several genes overlap with differentially abundant proteins listed above, including YWHAG, RAP1B, HAPLN2, ANK1, and CTSD. Although not all of the 58 labeled proteins were selected after the applied thresholds (Log 2-fold-change of 0.32 and p-value <0.05), abundances ratios in PD vs. Control and PD vs. PSP were calculated (Supp Fig. 4b). Among listed proteins, SOD1 and SLC16A1 had ratios > 1 in PD vs. Control and PD vs. PSP, suggesting these proteins are more abundant in PD bdEVs. On the other hand, alpha-synuclein (SNCA) had ratios < 1 in PD vs. Control and PD vs. PSP, suggesting lower abundance in PD, whereas Parkinson’s disease protein 7 (PARK7) returned divergent ratios of 0.87 and 1.103 in PD vs. Control and PD vs. PSP respectively, suggesting that PARK7 is less abundant in PD vs. control but more abundant in PD vs. PSP.

## DISCUSSION

### Summary of results

EV biomarkers of tissue pathology may be most valuable if they migrate into and can be found in accessible biofluids. To know what to look for in the periphery, we must thus know what is present in the cell and tissue of origin. Here, we report proteomic profiling of bdEVs purified from PD, PSP, and control brain tissues, finding several markers that differentiate groups. To the best of our knowledge, SLC8A2 and HAPLN2 differences are described for the first time as distinguishing bdEVs in PD. Some of these proteins, especially those implicated in endosomal pathway regulation, may serve not only as possible biomarkers but also as indicators of mechanisms in PD. Additionally, surface phenotyping of intact bdEVs showed the presence of selected neuronal, microglial, astrocytic, and endothelial markers that could be used in future to separate putative bdEVs from peripheral samples. We would now like to highlight several proteins of interest that we evaluated.

#### SLC8A2

SLC8A2 (NCX2) is a brain-specific Na+/Ca2+ exchanger (Jeon et al., 2003)(Calabrese et al., 2022). Mitochondrial SLC8A2 regulates calcium exchange in dopaminergic neurons, preventing calcium-dependent neurodegeneration (Gandhi et al., 2009; Wood-Kaczmar et al., 2008, 2013). NCX2 was reported to play a neuroprotective role in the ischemic model of the adult mouse brain (Jeon et al., 2008). Examination of NCX2 in synaptosomes of human brain specimens of Alzheimer’s disease revealed an increase of NCX2 in positive terminals and a colocalization with amyloid b in synaptic terminals (Sokolow et al., 2011). Because of its brain tissue specificity, localization at the cell membrane, presence in EVs, and potential involvement in PD, NCX2 is a promising candidate for more in-depth validation of mitochondrial dysfunctions seen in PD at the EV level.

#### HAPLN2

HAPLN2, also known as brain-derived link protein1 (Bral1), was more abundant in bdEVs in PD as previously reported in tissue (Liu et al., 2015). HAPLN2 is thought to be a secreted protein and to play a role in extracellular matrix formation and maintenance and neuronal conductivity (Bekku et al., 2009, 2010). Elevated HAPLN2 was linked to alpha-synuclein aggregation in an animal model of PD (Wang et al., 2016). More recently, HAPLN2 was reported (Teeple et al., 2021) to have significantly greater expression across oligodendrocyte clusters in PD vs. controls. Our results suggest the possibility that HAPLN2 overexpression is also reflected in EV association. We speculate that this protein might be involved in spread of PD pathology.

#### TMEM119

Microglially-enriched TMEM119 was elevated in PD and PSP in our assays. Although TMEM119 has been reported as specific for microglia, at least in the CNS (Bennett et al., 2016; Q. Li & Barres, 2018; Zhang et al., 2014), its levels may fluctuate during neuroinflammation (Bennett et al., 2016), and TMEM119-positive cells declined in a PD mouse model (George et al., 2019). We detected TMEM119 in some samples but not in others. Further studies are thus needed to determine if TMEM119 will be a valuable candidate for obtaining EVs from biofluids to assess the health of CNS microglia. Given the differing directions of regulation in neuroinflammation and PD, it may also be necessary to evaluate TMEM119 in patients at different stages and with different co-morbidities to assess its value in a diagnostic tool.

#### L1CAM

L1CAM affinity strategies have been used to obtain peripheral EVs of putative neuronal origin for biomarker discoveries, including in PD (Dutta et al., 2021; Jiang et al., 2021; Shi et al., 2014). However, neuronal specificity (Gutwein et al., 2003, 2005) and EV association (Norman et al., 2021) have both been questioned (Gomes & Witwer, 2022). Here, even in bdEVs, we detected only very low signals for L1CAM compared with other neuronally enriched markers, such as NCAM, NRCAM, and CD90 (Fig 3d). Like L1CAM, these proteins are also found on non-neuronal cells and outside the CNS.

In conclusion, multiple apparent PD markers were identified in this study, offering opportunities for follow-up. A strength of our study was the use of complementary profiling approaches. While the relatively unbiased proteomics approach was not specifically focused on membrane proteins (Hu et al., 2018), it was complemented by a targeted multiplexed ELISA approach to measure brain cell type-specific markers including membrane-associated proteins. Despite a relatively modest group size, significant differences were detected across categories. These differences are thus even more likely to be detected in larger validation cohorts. In sum, our study identified several brain cell type-specific markers on the surface of bdEVs that may be exploited for detection in the periphery, as well as markers that distinguish disease and control groups.

---

**Supp Fig 1.**
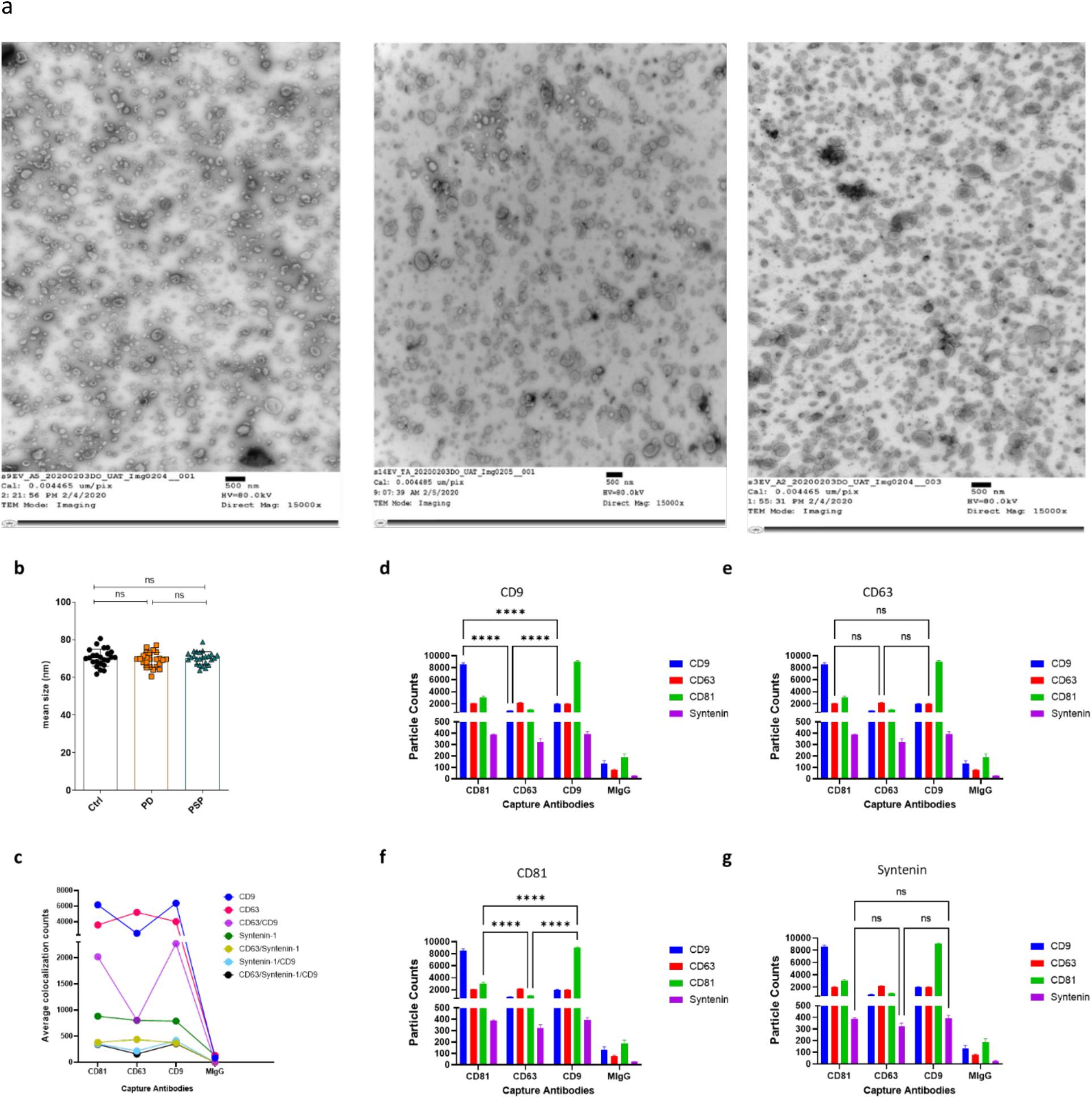
Minimal characterization of bdEVs. (a) Representative transmission electron micrographs (TEM. scale bar = 500 nm). (b) Mean diameter of human bdEVs as measured by nano-flow cytometry (NFCM) for control (black), PD (orange), and PSP (green) and normalized by tissue input. Data are presented as mean +/- SD. ns: no significant difference (P>0.05), Kruskal-Wallis ANOVA test. (c) Mean colocalization counts by SP-IRIS for each capture antibody for single fluorescently-labeled antibodies or combinations, as indicated: CD9, CD63, CD81, and syntenin. In (d-g), statistical analysis is presented for SP-IRIS-detected differences in fluorescence detection of CD9 (d), CD63 (e), CD81 (f), and syntenin (g) associated with each capture antibody, as indicated. Note that the same data are shown in each panel; the multiple panels are to highlight the different comparisons. Results are presented as mean +/- SD. **** = p < 0.0001. and ns = non-significant (two-way ANOVA) of three capture spots as indicated (CD81. CD63, CD9).

**Supp Fig 2.**
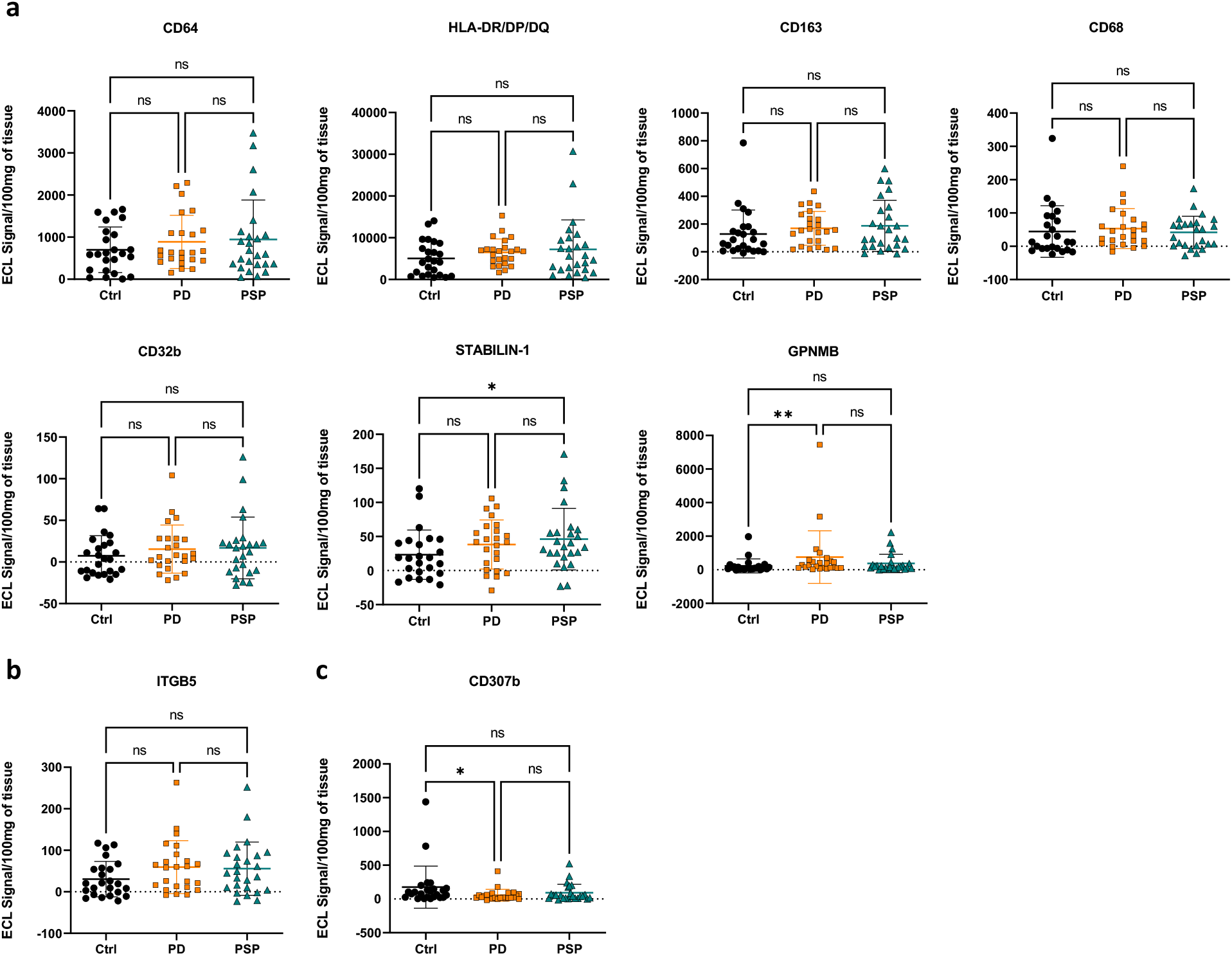
bdEV phenotyping by multiplexed ELISA. Comparison of control (Ctrl), PD, and PSP groups for (a) microglia, (b) microglia and astrocytes, and (c) microglia/neurons/astrocytes (c). Data were normalized per 100 mg of tissue input and reported as mean +/- SD. * = p < 0.05. ** = p < 0.01. and ns = non-significant (Kruskal-Wallis ANOVA).

**Supp Fig. 3:**
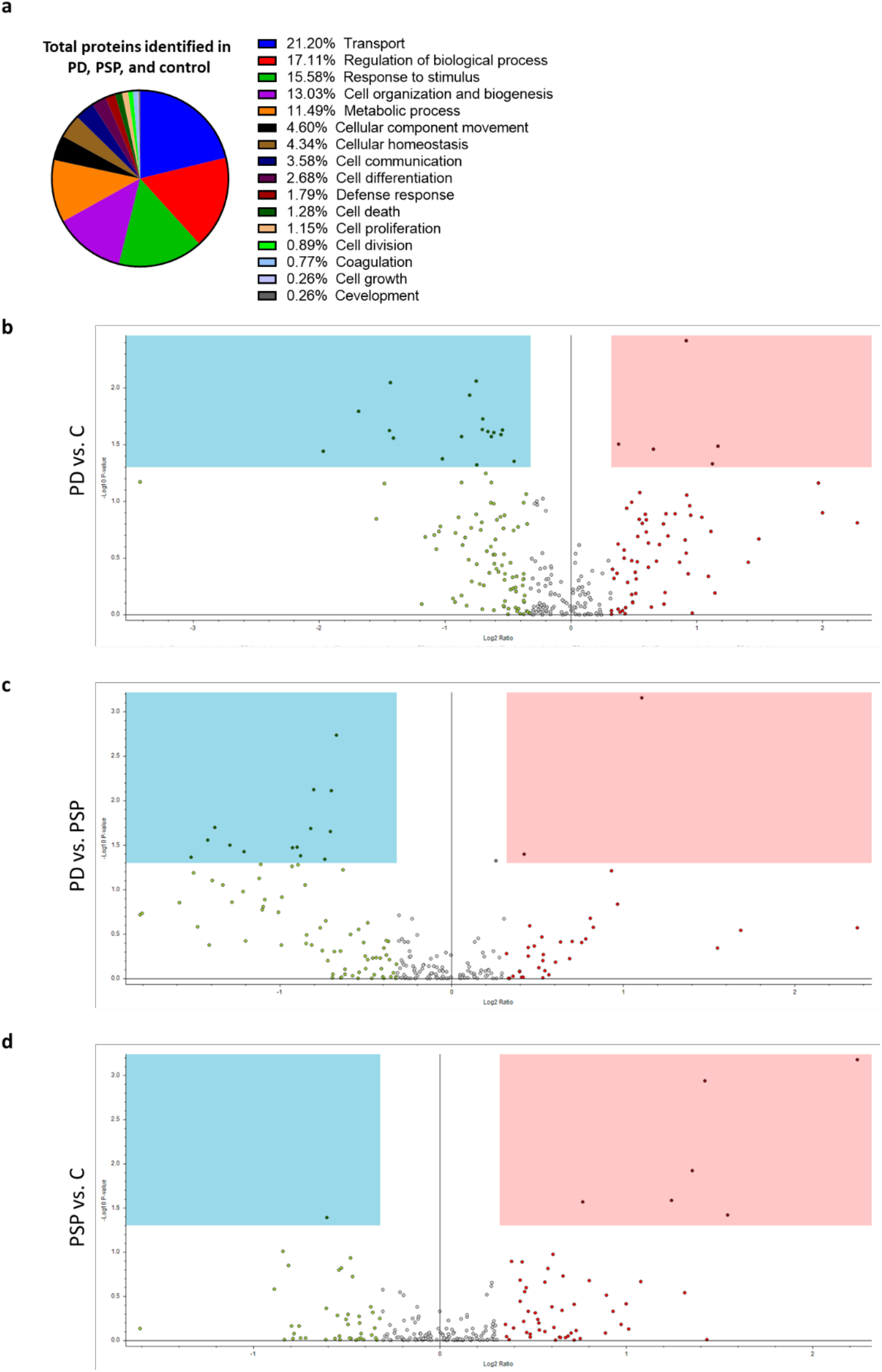
Proteomic profile of bdEVs identified in PD, PSP, and control. a) Gene Ontology terms associated with the proteome of bdEVs. b-d) Volcano plots of individual proteins of PD vs. control (b), PD vs. PSP (c), and PSP vs. control (d). Light blue squares: considered significantly more abundant. Light red squares: considered significantly less abundant.

**Supp Fig. 4:**
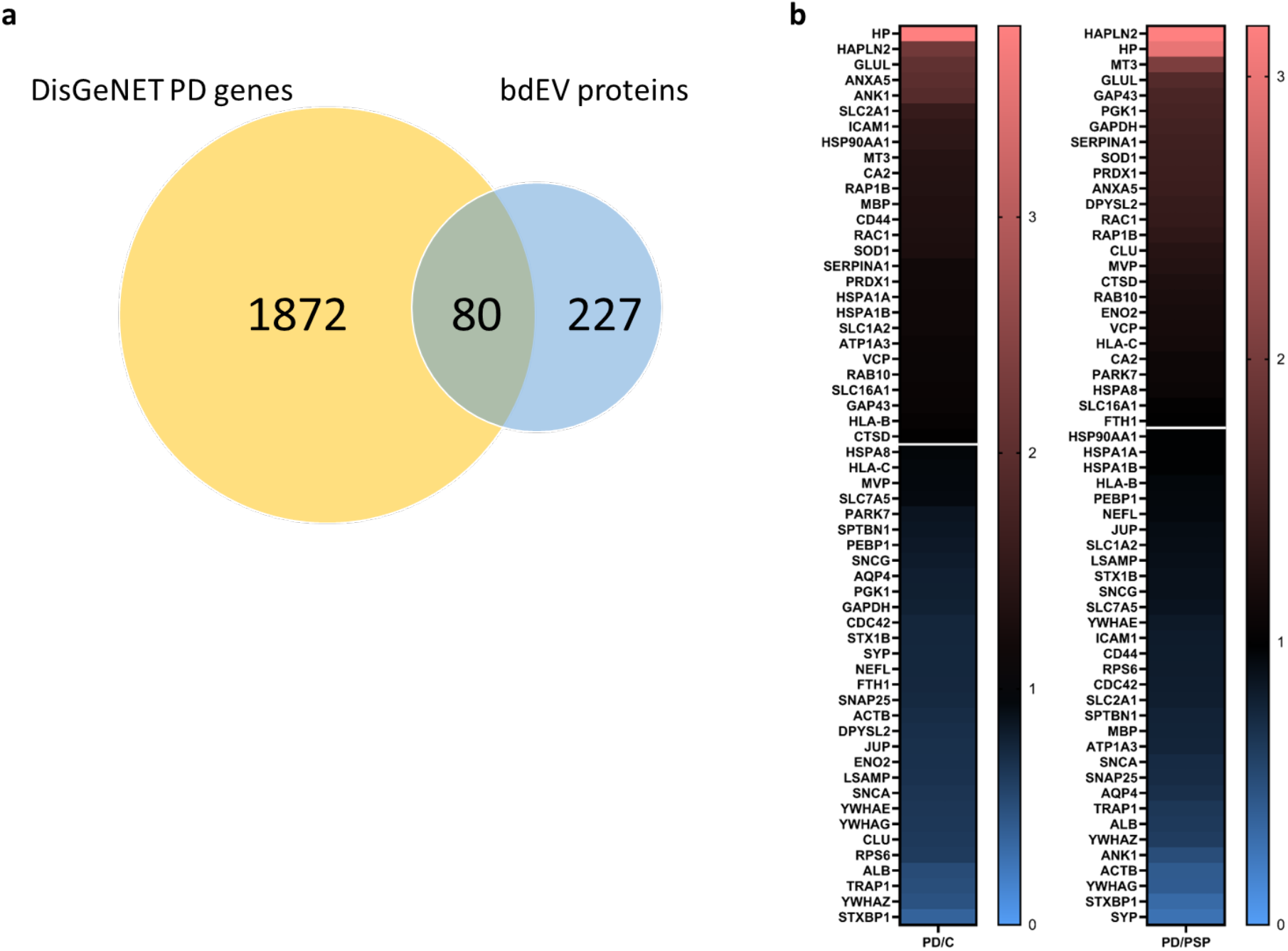
PD-related proteins and the bdEV proteome. a) bdEV overlap: PD-related proteins listed in the DisGeNet database and proteins detected in the bdEV proteome and identified in the three tested groups. b) Abundance ratios of individual bdEV proteins from average biological replicates: PD/Control (left) and PD/PSP (right) per the scale shown on the right. White lines correspond to ratio=1.

## Notes

### Competing Interest Statement

RN, EG, and DAR are employees of Meso Scale Diagnostics, LLC. KWW privately consults in the extracellular vesicle space and has received stock options from NeuroDex, Inc., who had no role in the design or interpretation of this study.

